# Synthesizer: Expediting synthesis studies from context-free data with natural language processing

**DOI:** 10.1101/053629

**Authors:** Lisa Gandy, Jordan Gumm, Benjamin Fertig, Michael J. Kennish, Sameer Chavan, Ann Thessen, Luigi Marchionni, Xiaoxan Xia, Shambhavi Shankrit, Elana J Fertig

## Abstract

Today’s low cost digital data provides unprecedented opportunities for scientific discovery from synthesis studies. For example, the medical field is revolutionizing patient care by creating personalized treatment plans based upon mining electronic medical records, imaging, and genomics data. Standardized annotations are essential to subsequent analyses for synthesis studies. However, accurately combining records from diverse studies requires tedious and error-prone human curation, posing a significant barrier to synthesis studies. We propose a novel natural language processing (NLP) algorithm, Synthesize, to merge data annotations automatically. Application to patient characteristics for diverse human cancers and ecological datasets demonstrates the accuracy of Synthesize in diverse scientific disciplines. This NLP approach is implemented in an open-source software package, Synthesizer. Synthesizer is a generalized, user-friendly system for error-free data merging.

## 1 Introduction

Today’s digital age has provided unprecedented access to high quality datasets collected from independent studies for every scientific discipline. Meta analyses that combine data across studies can provide comprehensive databases to elucidate novel scientific conclusions. For example, in the case of human cancer public domain databases such as Gene Expression Omnibus (GEO) and Array Express have gathered high-throughput, multi platform characterization of tumors from thousands of independent datasets (see Figure 1a for numbers of datasets per database). Combining data in public databases also greatly enhances the number of tumors available for molecular profiling by any individual investigative team [12] [5] enabling public domain analysis tools sufficient sample size to validate molecular biomarkers [1, 11].

**Figure 1:**
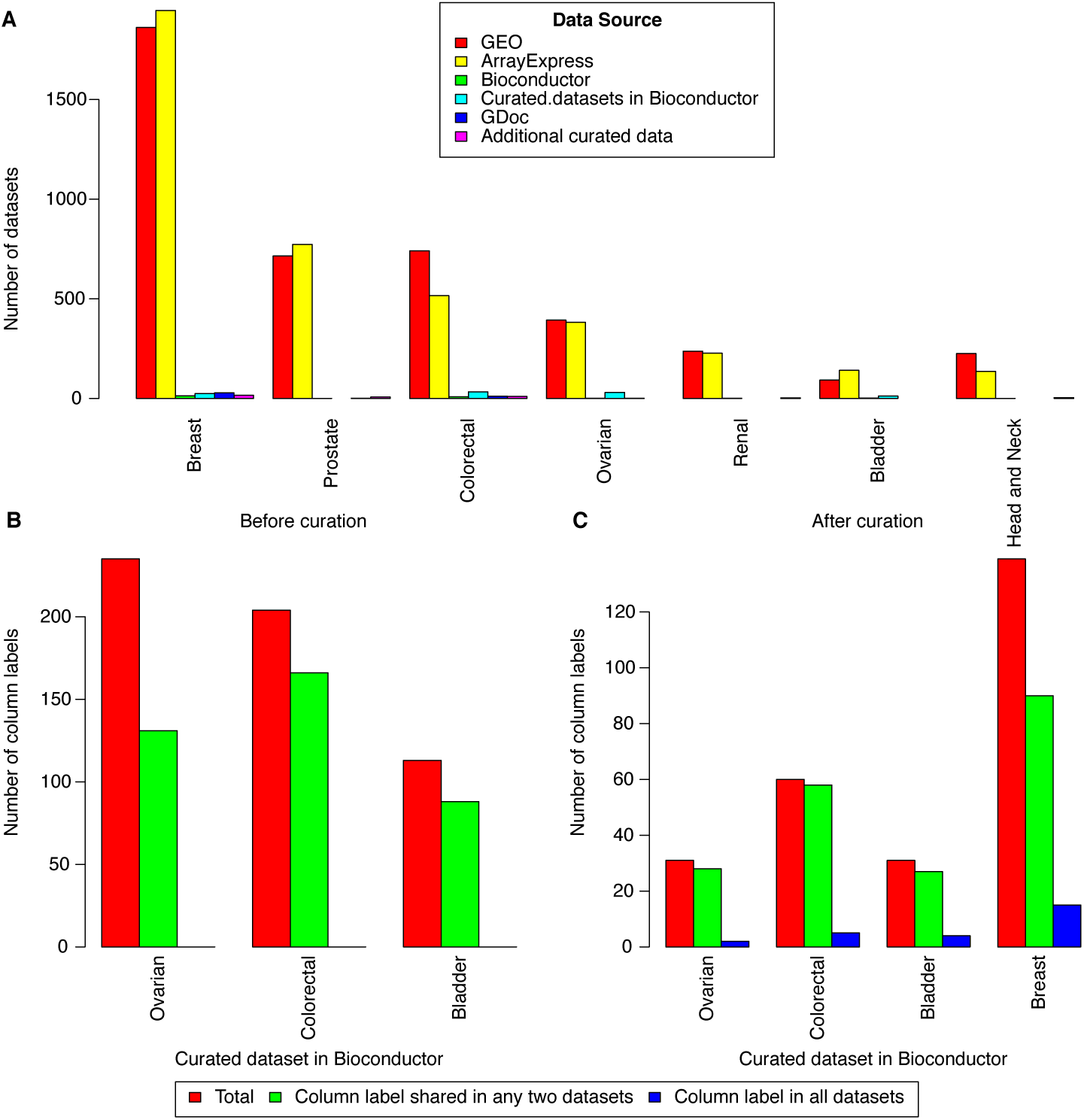
Number of datasets available before and after curation.

Though using synthesis studies does mitigate the cost of data collection, there are still costs associated with using disparate datasets. A critical and time intensive first step in using multiple datasets is to merge the annotations of each sample manually. These sample annotations are frequently provided as labels in data tables determined by an individual investigator that do not conform to any standards. For example, in a set of curated genomic databases, none of the column names for sample phenotypes were shared across every dataset prior to curation (Figure 1b). Only 25% of the sample annotations were shared across all these datasets after intensive human curation to match sample phenotypes to a common standard (Figure 1c). In regards to cancer data, there are thousands of genomic datasets in the public domain, however only a handful of the datasets feature curated clinical annotations (as illustrated in Figure 1a) due to the time cost associated with manual annotation [10, 13].

Prior work has been done in Natural Language Processing (NLP) and computational linguistics to map datasets to existing ontologies in an effort to aid scientists when synthesizing datasets. Numerous systems, including notably MetaMap [2] and Mgrep [6] have been developed to link biomedical text to existing terms and index biomedical literature. These programs are linguistically sophisticated, employing word sense disambiguation, text negation, and detects author-defined abbreviations and acronyms. Shah et al. apply MetaMap and Mgrep to obtain ontology based labels of genomics data. This work has been applied to select appropriate datasets to analyze in order to explore the interaction between phenotype, disease, environmental and experimental data [4] and impute phenotypes [18]. Nonetheless, robust statistical analysis of terms that are not represented in standardized ontologies or comparisons of classes of samples within an ontology still requires substantial manual standardization of the sample annotations provided in unformatted data tables.

To address this limitation, the field of NLP is advancing in its ability to recognize commonalities between words. In their infancy, NLP and information retrieval (IR) systems related words by querying for exact matches in sets of documents. Today’s NLP and IR systems are more advanced, and can define a “semantic context” that describes related words. These contexts are frequently defined from synonyms, hypernyms, and collocates between words in independent, lexical databases like WordNet [8] or the Corpus of Contemporary American English (COCA) [7]. Diverse applications built on NLP concepts such as Google, metaphor identification software [9], and sentiment analysis [3] all use semantic contexts. We propose that the ability to integrate text data without an explicit context can be used to annotate samples across a wide range of biomedical studies. To our knowledge, there is currently no publicly accessible software that automatically combines unformatted data tables from disparate datasets using NLP concepts. The closest application in terms of functionality was Google Refine. The Refine software allowed the user to import data in various formats and did automatically combine identically labeled columns. However, unlike the software presented in this paper, Google Refine did not automatically combine data with differently named columns. In addition Google Refine has been renamed to OpenRefine [17], and is no longer supported by Google.

Therefore, we propose a novel natural language processing algorithm to mine and standardize data tables by introducing semantic context. This NLP algorithm, combined with a user-friendly drag and drop web application called Synthesizer facilitates the seamless standardization of data. We demonstrate the efficacy of the algorithm and software to standardize patient phenotype annotation using a large collection of genomics datasets for four human cancers. Only one of the cancer data sets (HNSCC) was used to tune the Synthesize algorithm, with all other data sets used for testing. Although trained on the HNSCC cancer dataset, the algorithm is generally applicable to standardize sample annotations for datasets in other disciplines, with demonstrated accuracy standardizing annotations in three ecological datasets.

## 2 Results

### 2.1 The Synthesize algorithm

We developed a novel natural language processing algorithm to standardize sample annotations called Synthesize (workflow outlined in Figure 2). The input of the algorithm is unformatted data, with column headers represented as variables and the variable values being the sample data given in the rows from the dataset (Figure 3a). The algorithm merges labels of sample annotations based on their similarity in a “semantic space”. Specifically, the algorithm queries the COCA database to find word collocates for each of the terms in a given column of data. An example of the semantic space generated in relation to candidate columns “gender” and “sex” is given in Figure 3b.

**Figure 2:**
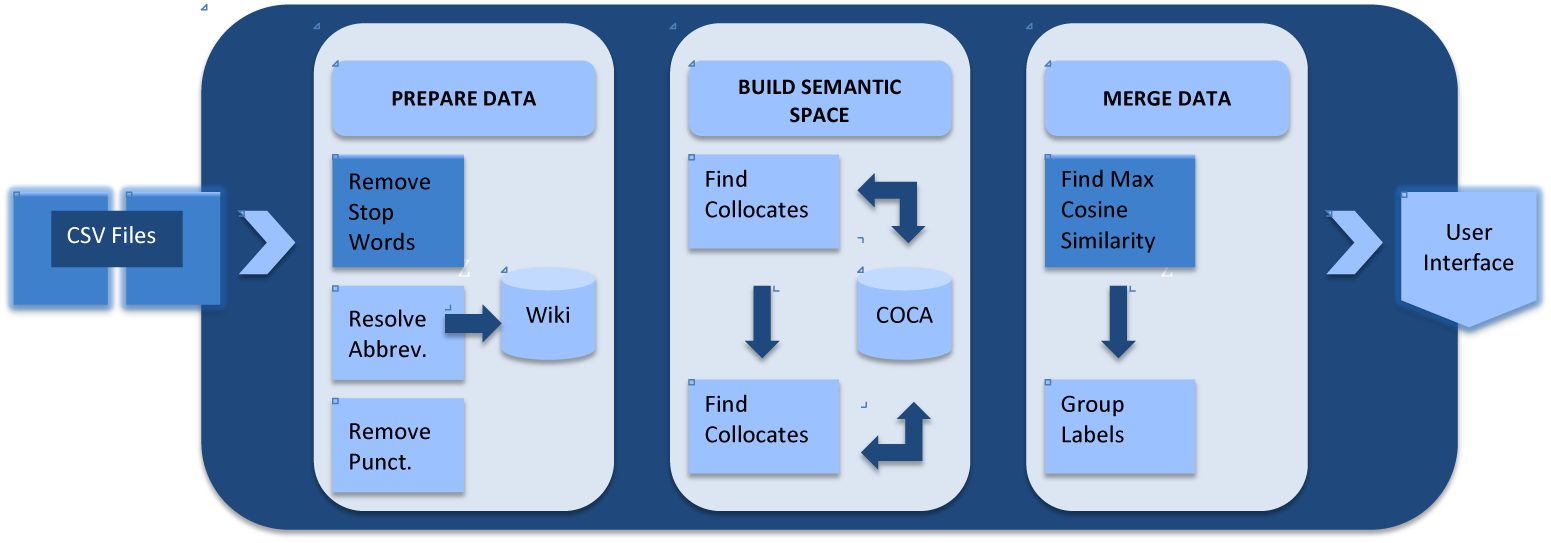
System figure for the Sythesizer application.

**Figure 3:**
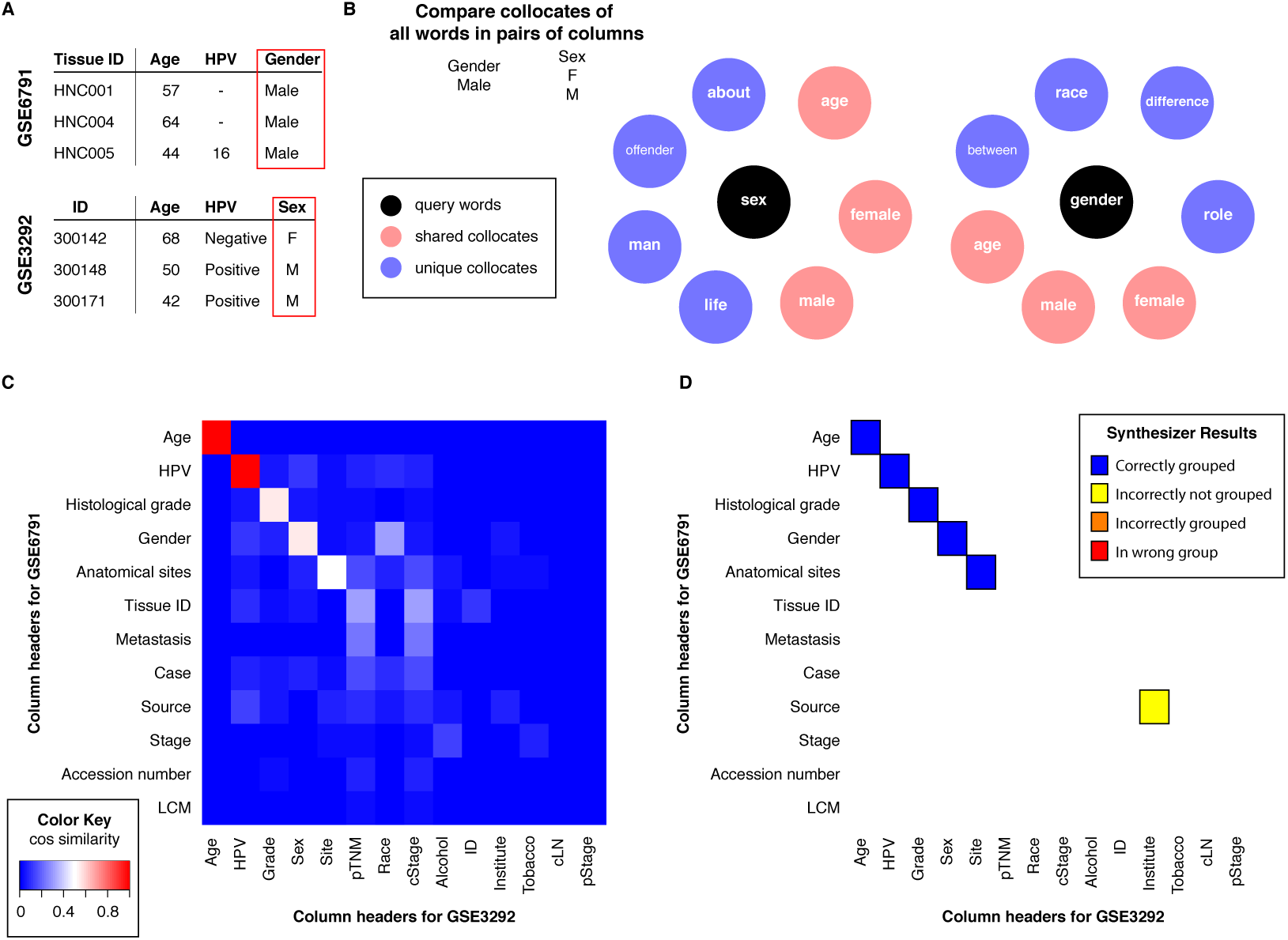
Synthesize algorithm overview.

A similarity score is computed for each candidate pair of columns to be merged. The similarity score is calculated by comparing the column header values and collocates of both for each pair. Two columns are marked as candidates to merge if their similarity score is above a threshold value and is higher than all other candidate pairs. We selected the threshold value to minimize standardization errors for annotations of genomics data from two head and neck [14, 16] cancer datasets. The resulting algorithm had only one classification error, missing standardization of the columns labeled “source” and “institute”. These columns contain abbreviated university names, which are not recognized as terms in COCA and therefore cannot be merged by our system (Figure 3d). The full Synthesize algorithm can be found in the appendix under Section 1: Synthesize Natural Language Algorithm. The link to the full Synthesize code, which is currently implemented in Python, can be found in Section 3 of the Appendix.

### 2.2 Dataset Description

In order to test the accuracy of the Synthesize algorithm we collected four independent datasets of cancer annotations from gene expression analysis. These cancer datasets were retrieved from Gene Expression Omnibus. For each cancer type considered, only datasets corresponding to clinical cohorts and with at least 10 samples were included. Studies for which no clinical or pathological information was available were excluded. For each dataset, phenotypic information was extracted from the ExpressionSet instance, saved in tabular format (TAB-delimited text) and manually curated/reviewed for data consistency and quality. Whenever possible column headers were relabeled based on the information contained in the column itself.

To assess the applicability of the Synthesize algorithm to merge annotations for synthesis studies in scientific disciplines beyond oncology, we tested the algorithm using three Ecology datasets. The datasets included data from seagrass monitoring, insect abundance and amphibian life history. In regards to the seagrass monitoring data, samples of eelgrass (Zostera marina L.) and widgeon grass (Ruppia maritima) were collected from Barnegat Bay and Little Egg Harbor Estuary in New Jersey along a gradient of human population density and development. Quadrant, core, and hand sampling of seagrass, following SeagrassNet monitoring and sampling protocols [15] were conducted between 2004 and 2013.

The insect abundance data set is a series of observations of specific lepidopteran taxa made by citizen science programs in North America. The numeric abundance is reported along with a taxon name, location, time, and some environmental measurements. The data are collected under a moderately standardized protocol, but several important differences exist in the terms used. The amphibian data set is the result of a literature search for extrinsic and intrinsic traits of Mexican amphibians. Data were manually entered into a spreadsheet by several different researchers.

### 2.3 Accuracy of Synthesize on merging annotations for cancer genomics datasets

We compare the accuracy of the columns in the cancer datasets combined with Synthesize to hand curation of the annotations. In Figure 4a we give a breakdown of error types. Errors can be broken down in three distinct categories: (1) not grouped by synthesis but grouped in hand curation (incorrectly not grouped), (2) grouped by synthesis but not in hand curation (incorrectly grouped), and (3) placed in different groups by synthesis and hand curation (wrong group). We find that Synthesize accurately merges 89% of 200 columns in 16 tables for breast cancer, 91% of 137 for colorectal cancer, 92% of 74 columns in 10 tables for prostate cancer and 100% of the 23 columns in 3 tables for renal cancer. As the number of overall columns increases the accuracy generally decreases but is still high.

**Figure 4:**
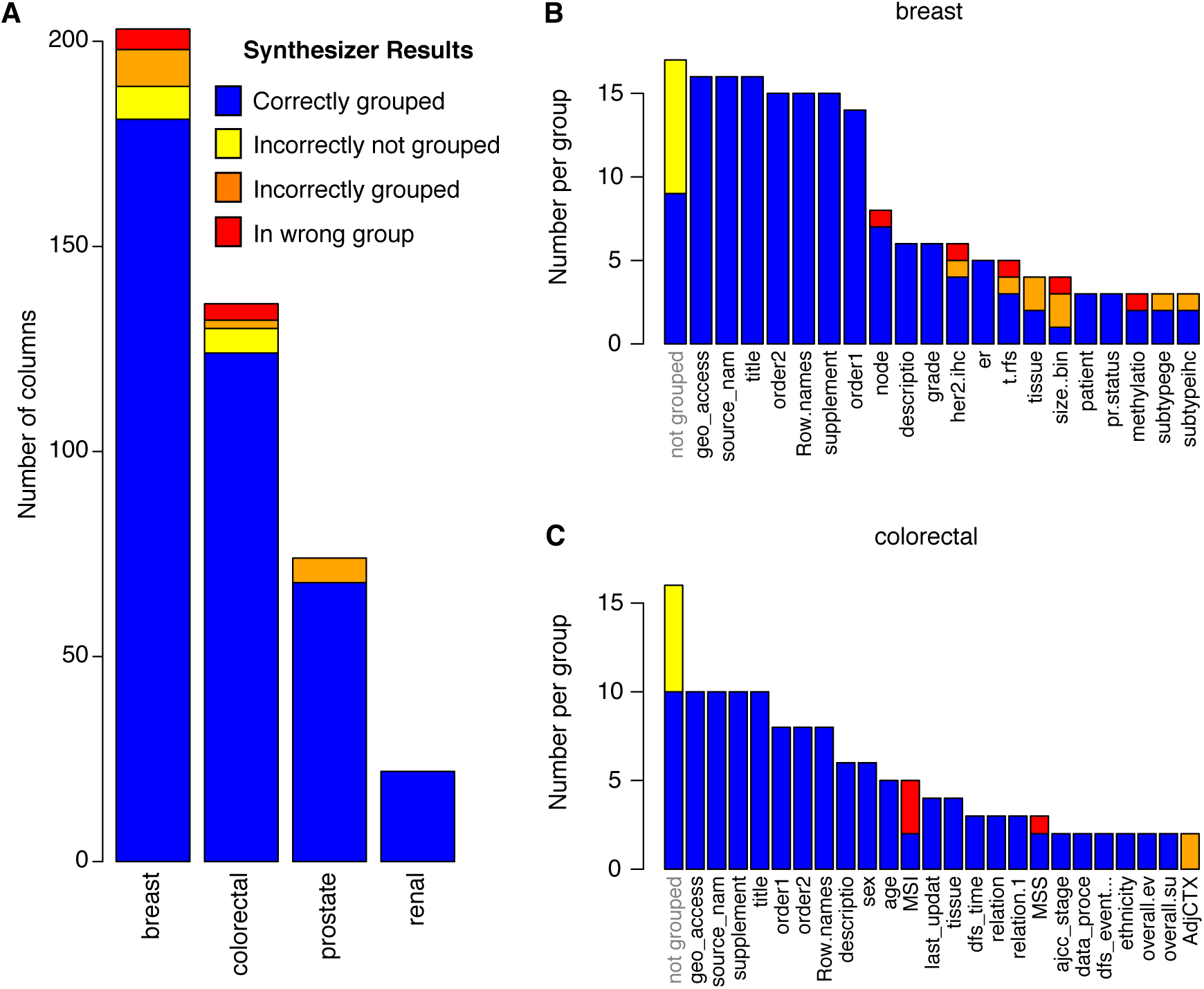
Synthesize accuracy with regards to cancer genomics datasets.

In Figures 4b and 4c we give a further breakdown of the two cancer datasets with the most errors: breast and colorectal. In these datasets, the most frequent errors of the three error types, 3% were in category 1, 5% in category 2 and 2% in category 3, meaning that the most errors occur when columns are grouped together when they should be left ungrouped (5% of the total number of annotations) as opposed to being incorrectly ungrouped (3%), or placed in the wrong group (2%). Within the breast cancer data, we observe errors merging the column labeled “pr” or “pr.status” in datasets GSE11001, GSE23593 with the “progesterone receptor status” in dataset GSE36774. In addition, the system was unable to merge the measurements for the HER2 receptor for a dataset that contained independent measurements with fluorescence in situ hybridization (column labeled “her.fish” in dataset GSE29431) and immunohistochemistry (column labeled “her.ihc” in dataset GSE29431). Finally, the system merged a column labeled “description.1” in the GSE23593 datasets to “methylation.barcode” in the GSE20711 and GSE20712 datasets due to the alphanumeric categories in both.

### 2.4 Accuracy of Synthesize on merging annotations for ecological datasets

In regards to the ecological data, the accuracy of the system compared to hand annotation was 85% of 410 columns in 9 spreadsheets correctly merged in regards to seagrass ecology data, 95% of 34 columns for insect data and 92% of 54 columns in regards to amphibian data. As with the cancer data, the accuracy decreases with the number of columns (see Figure 5a). Figures 5b–d, summarize error types for each ecology dataset. For these data, 2% of the annotations were incorrectly not grouped, 2% grouped when they should be ungrouped, and 10% placed in the wrong group. As with the cancer datasets, as the number of overall columns increases the accuracy generally decreases but is also still high.

**Figure 5:**
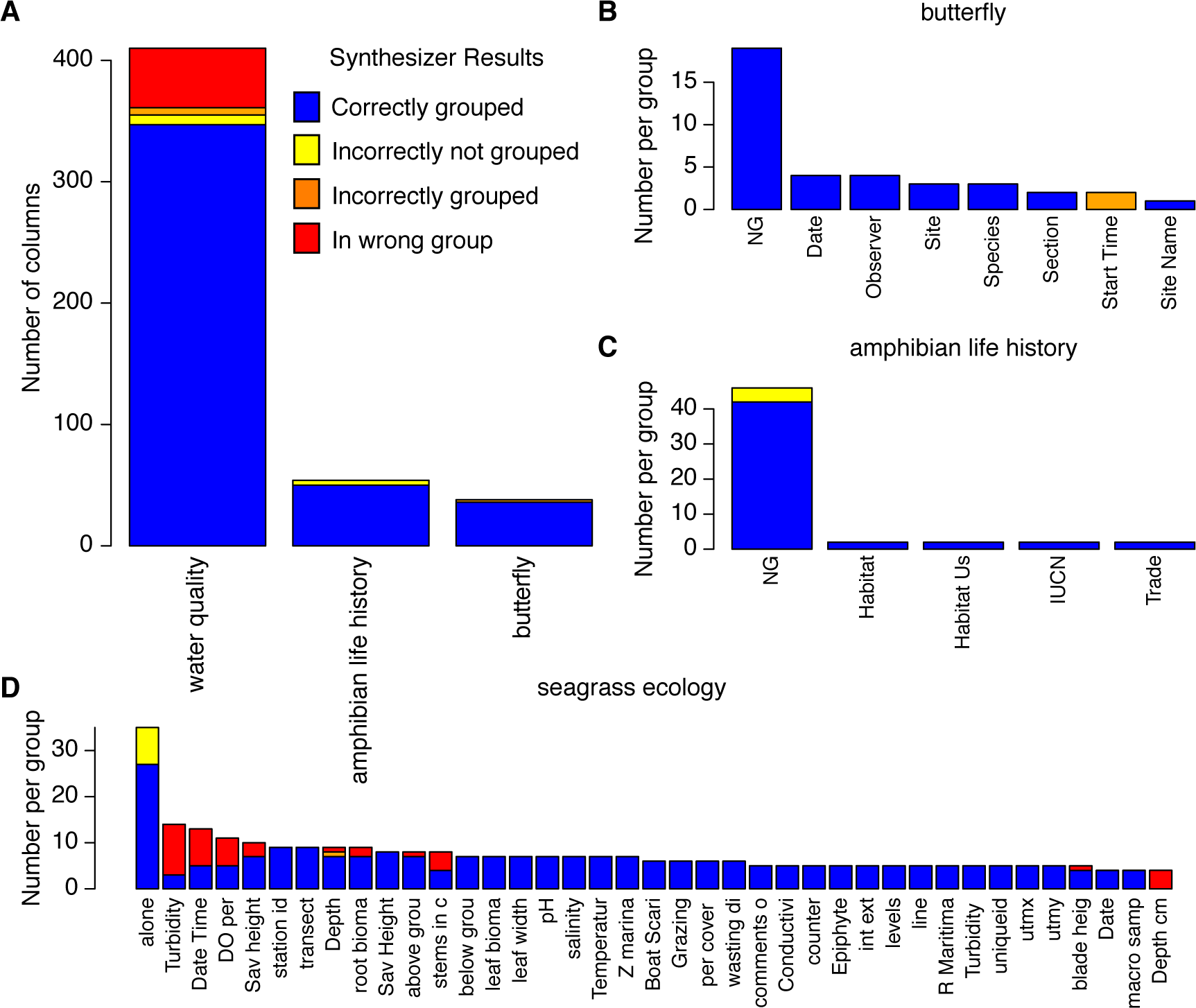
Synthesize accuracy with regards to ecology datasets.

### 2.5 Synthesizer interactive annotation merger software

The Synthesizer system is designed to implement the Synthesize algorithm in an automated, flexible web based system. This system uses a graphical user interface that employs point and click to upload files, merge labels from sample datasets and download merged datasets in a seamless workflow. The user begins with uploading sample datasets using a simple drag and drop interface. The users drag their spreadsheets into the html interface. Alternatively, they can upload the data by clicking and using the native file browser system for their operating system to find their files and submit them. The system then uploads the data to our local data server, and implements the Synthesize algorithm to merge sample annotations.

As shown below in Figure 6, the Synthesizer interface provides suggested labels on the left hand side of the screen. The user now has the option to click and move labels between merged groups, and also create new groupings. The interface also features the ability to change dataset values individually or by using a global search and replace option. Once the user has merged labels and changed the data to their satisfaction, they can then obtain a new merged spreadsheet by choosing the merge option provided in the interface. The human-computer interaction implemented in this software enables users to correct for all errors introduced by the NLP algorithm. As a result, it is possible to obtain perfect accuracy in regards to merging data but with the added benefit of far less work when manually merging all data. Further description of the graphical user interface software can be found in Section 2 of the appendix and a link to the public access code (written in Cappucino) can be found in Section 3 of the appendix.

**Figure 6:**
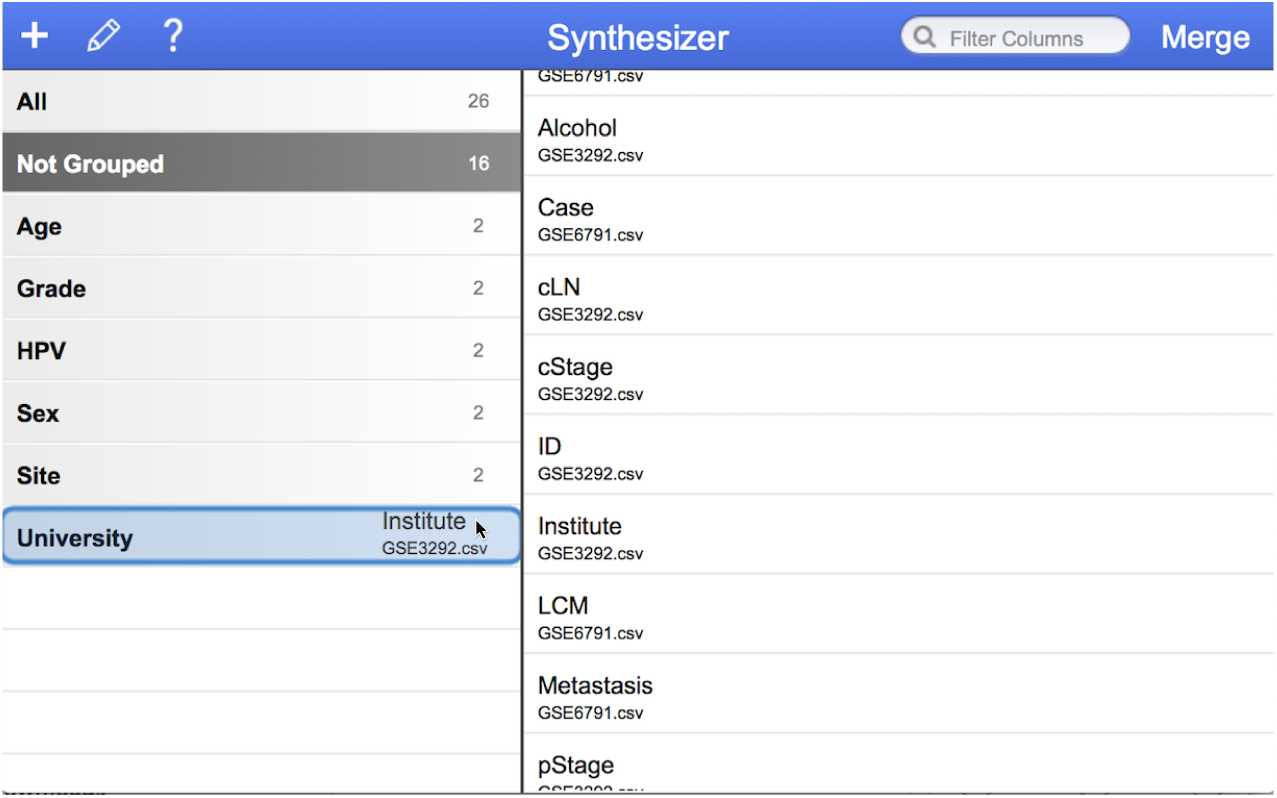
Synthesizer graphical user interface.

## 3 Discussion

This paper presents Synthesizer, which eases the time challenges related to synthesis studies through the ability to quickly and easily combine data. We develop a new NLP algorithm, Synthesize, to mere sample annotations, with an intuitive interface for human-computer interactions to refine merged columns in data. We train the Synthesize algorithm on annotations for head and neck cancer genomics datasets, and demonstrate that the algorithm retains high accuracy (ranging from 89%-100%) merging the annotations for independent cancer genomics datasets. The overall accuracy for ecology data (ranging from 85%-95%) was comparable to that of the cancer data, despite the fact the system was trained on annotations from head and neck cancer and abbreviations were resolved against medical terms. In all cases, the number of errors scaled with the total number of columns in the datasets. For example, the seagrass ecology dataset had nearly double the number of columns (410) of the largest cancer dataset (200) and a corresponding decrease in accuracy (85% for seagrass data and 89% for breast cancer). Therefore, we anticipate that Synthesizer would have similar accuracy in merging annotations from additional medical and scientific disciplines. The flexible interface for Synthesizer to input unformatted, comma delimited text files of data further supports data merging for such cross-disciplinary synthesis analysis. In all cases, we found that errors in merging annotations with Synthesizer were attributed to three primary factors: (1) use of abbreviations, (2) multiple columns in one dataset that described a measurement, and (3) bias in the NLP algorithm towards grouping sample labels.

The first factor involves the incorrect resolution of abbreviations by the system. Our system currently uses a generalized list of medical abbreviations. We chose a generalized list so that the system could be more adaptable to different types of datasets. However consider the case of a column labeled “pr” in the breast cancer data referring to the status of the “progesterone receptor” in a breast tumor. In this case the system would resolve “pr.status” to “prothrombin ratio status” and therefore would be less likely to merge “pr.status” and another column titled “progesterone receptor”. Such errors could be mitigated in future extensions of Synthesizer that enable the user to specify a context-dependent dictionary to resolve abbreviations.

The second factor regarding incorrect grouping is the different ways in which data is divided into columns per dataset. For instance in the breast cancer dataset, one spreadsheet features a column labeled “her” referring to the presence of the genomic amplification of the HER2 gene. In another spreadsheet this HER2 status is measured with two independent assays by immunohistochemistry (her.ihc) and fluorescence in situ hybridication (her.fish). In this case we have a missing metadata issue, as it is unclear even to a human expert in genomic testing how these columns should be combined without additional information. The current system groups “her” and “her.ihc” together, which is arguably correct, though her.fish is not represented in the merger in this case.

Finally, the system is biased towards grouping data, resulting in liberal sample mergers. An example would be a column labeled “description.1” in the breast cancer dataset, which contains information regarding a phenotype encoded using letter and number combinations such as A1, B1,… F1. In the system this column is grouped with “methylation.barcode” from another spreadsheet in the same dataset. Although incorrect, this merge occurred because the methylation barcode column has a series of letters then numbers just like description.1.

In terms of work with medical ontologies, we see this work as a complement and not a de facto replacement. One could imagine that in the future Synthesizer could offer a list of widely used ontologies in the medical field and then aid the user in mapping their current dataset using the ontology. In addition, Synthesizer could be used to help researchers map two ontologies onto each other. In future enhancements of the system, this mapping could be saved and used for future merges of datasets. By incorporating medical ontologies, Synthesizer would encourage users to make use of standardized mappings, but with the aid of its automated merge capabilities and easy to use interface.

## 4 Acknowledgments

Elana Fertig was supported by the National Institutes of Health (NIH) National Cancer Institute (NCI) (UL1 TR 001079). Luigi Marchionni was supported by the National Institutes of Health (NIH) National Cancer Institute (NCI) (P30 CA006973 and UL1 TR 001079), and by the Cleveland Foundation The Helen Masenhimer Fellowship Award. Seagrass data from the Barnegat Bay-Little Egg Harbor Estuary are the result of research sponsored by the New Jersey Sea Grant Consortium (NJSGC) with funds from the National Oceanic and Atmospheric Administration (NOAA) Office of Sea Grant, U.S. Department of Commerce, under NOAA grant number (NA10OAR4170075) and the NJSGC.

